# Heart rate persistence index (HRPI): a threshold-free wearable metric for sustained HR elevation

**DOI:** 10.64898/2026.04.07.716987

**Authors:** Ren Zhang

## Abstract

Wearable devices generate dense longitudinal heart rate (HR) data, but summarizing sustained heart rate elevation in a single daily metric remains challenging. We developed the heart rate persistence index (HRPI), defined as the largest integer *k* such that at least *k* minutes in a day have HR ≥ *k* bpm. For example, an HRPI of 105 means daily HR was ≥105 bpm for at least 105 minutes. HRPI is threshold-free and integrates magnitude and duration of elevated HR into a single interpretable value. Using multi-day wearable recordings from a PhysioNet dataset, we show that HRPI captures structure beyond mean HR, reflects variability-related features, and exhibits robust day-to-day stability. In an independent healthy cohort, HRPI declines strongly with age, supporting physiological relevance. HRPI offers a compact, interpretable, and robust summary of sustained HR elevation for longitudinal wearable studies, providing information easily accessible to both specialists and nonspecialists.

## Introduction

Wearable devices now generate continuous physiological data at unprecedented scale [1]. Heart rate (HR) is among the most accessible and widely used of these signals, reflecting autonomic state, physical activity, fitness, cardiometabolic health, and disease risk [2-4]. In such longitudinal data, both the magnitude and duration of HR elevation are important, because transient spikes differ fundamentally from sustained high HR in both physiological meaning and potential clinical relevance [4]. Existing daily HR summaries each capture only part of the information contained in continuous HR data. Mean HR reflects overall level but does not distinguish transient from sustained elevation. Maximum HR is dominated by brief peaks. Threshold-based measures, such as minutes with HR >100 bpm, are intuitive but rely on arbitrary cutoffs. Percentile-based metrics characterize the upper portion of the daily HR distribution, yet they do not directly unify magnitude and duration in a single interpretable quantity. Consequently, there remains a need for a simple daily summary that is threshold-free, parameter-free, intuitive, and capable of capturing the persistence of elevated HR in a single number.

To address this gap, we developed the heart rate persistence index (HRPI), a new metric for summarizing sustained daily HR elevation. HRPI is designed to provide a compact and interpretable one-number summary of persistent high HR without relying on arbitrary thresholds or tunable parameters. By jointly incorporating magnitude and duration, HRPI captures two-dimensional information in a simple form and reflects a distinct aspect of daily HR dynamics. Such a metric can be useful from both biological and practical perspectives. Biologically, sustained HR elevation may capture information not contained in mean HR alone, including the distribution and persistence of elevated HR values across the day. Practically, a simple and interpretable daily summary of persistent HR elevation could be valuable for wearable-based longitudinal monitoring, phenotyping of HR patterns, and potential clinical or behavioral applications.

In this study, we developed HRPI and evaluated it in two complementary datasets. Using a wearable dataset with repeated daily measurements, we examined how HRPI relates to conventional HR metrics, tested whether it captures structure beyond mean HR, and assessed its day-to-day stability within individuals. Using an independent healthy reference cohort spanning infancy through adulthood, we then examined whether HRPI shows a physiologically meaningful association with age.

## Materials and Methods

### Data sources and processing

Continuous wearable data were obtained from the PhysioNet “BIG IDEAs Glycemic Variability and Wearable Device” dataset, comprising multi-day recordings from 16 participants using an Empatica E4 device [5-7]. HR data were aggregated to minute-level resolution. Daily segments were defined as 24-hour periods, and days with insufficient coverage were excluded based on data completeness criteria.

Age-dependent analyses were performed using the PhysioNet dataset “RR interval time series from healthy subjects” [7-9]. This dataset contains 24-hour RR interval recordings from healthy individuals across a broad age range, with age information provided in the accompanying metadata. RR intervals (ms) were converted to instantaneous HR as HR (bpm) = 60000 / RR (ms) and aggregated to minute-level resolution. Subject-level HRPI values were then computed.

### Definition of HRPI

For a given day, let *h*_(1)_ ≥ *h*_(2)_ ≥ … ≥ *h*_(n)_ denote the minute-level HR values sorted in descending order. HRPI was defined as

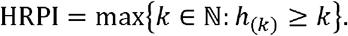

Thus, HRPI represents the largest integer *k* for which at least *k* minutes in that day had HR ≥ *k* bpm.

### Data processing, metric computation, and statistical analysis

For each subject-day, we calculated mean HR, 90th percentile HR (p90 HR), 95th percentile HR (p95 HR), standard deviation of HR (SD HR), coefficient of variation of HR (CV HR = SD/mean), and minutes with HR >100 bpm. Associations between HRPI and conventional HR metrics were assessed using Pearson correlation. To isolate the component of HRPI independent of mean HR, linear regression was performed with HRPI as the dependent variable and mean HR as the predictor, and residual HRPI values were extracted. Within-subject stability was quantified as the coefficient of variation across days for each metric. For age analysis, subject-level HRPI values were regressed against age, with age displayed on a logarithmic scale for visualization. All analyses were performed in Python using standard scientific libraries. All statistical tests were two-sided, and *p <* 0.05 was considered statistically significant.

## Results

### Geometric definition of HRPI

To capture the sustained elevation of HR throughout a day, we developed the HRPI, which is a rank-based measure that integrates the magnitude and duration of elevated HR within a day. Minute-level HR values are sorted in descending order and plotted against cumulative time. In this representation, HRPI is the largest integer *k* such that at least *k* minutes have HR ≥ *k* bpm, corresponding to the intersection of the ranked curve with the identity line (*y = x*) (Fig. 1A). This geometric formulation gives HRPI a straightforward interpretation: it reflects sustained elevation of HR rather than isolated peaks or average levels. Higher HRPI values occur when elevated HR is maintained for longer durations, shifting the ranked curve upward and extending its intersection with the identity line.

**Fig. 1.**
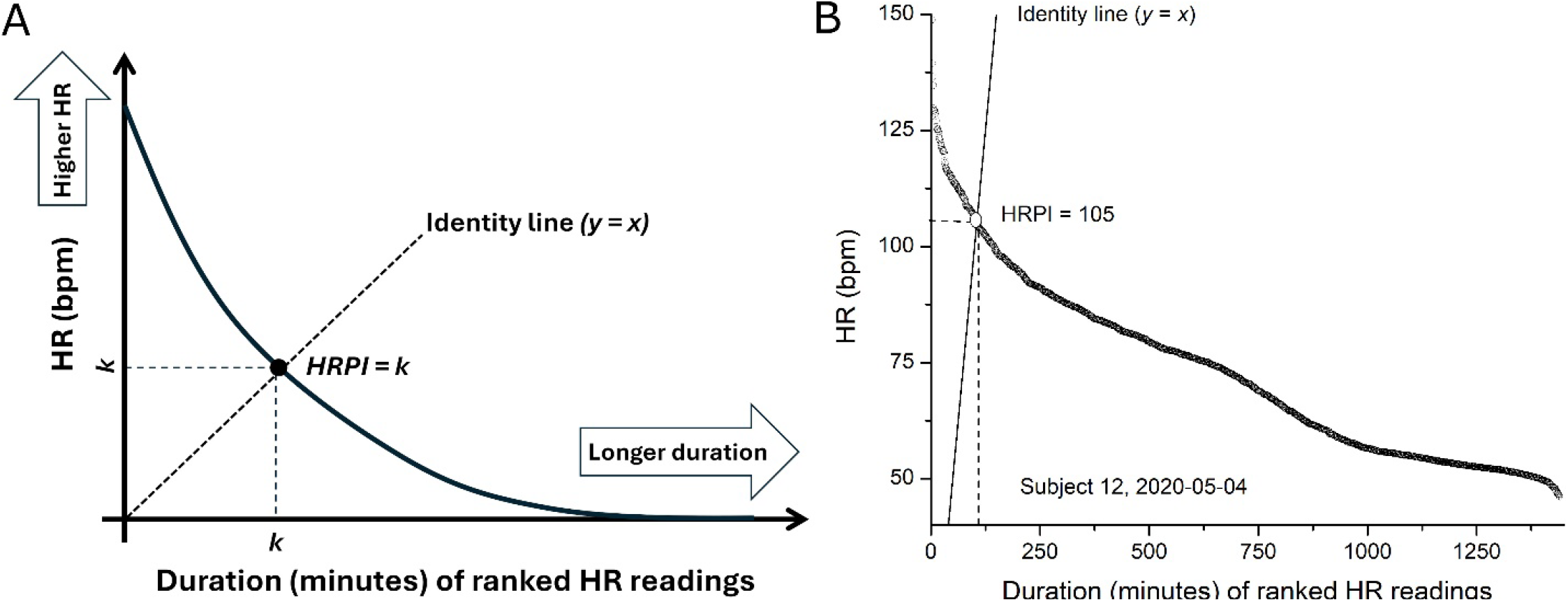
Definition of HRPI. A) Geometric representation of HRPI. Minute-level HR values within a day are ranked in descending order and plotted against cumulative duration. HRPI is the largest integer *k* such that at least *k* minutes have HR ≥ *k* bpm, corresponding to the intersection of the ranked curve with the identity line (*y = x*). B) Example from a representative subject-day. Minute-level HR values are ranked in descending order. In this example, HRPI = 105, indicating 105 minutes with HR ≥ 105 bpm.

To illustrate the definition using subject-level wearable data, we used the PhysioNet “BIG IDEAs Glycemic Variability and Wearable Device” dataset, which comprises multi-day Empatica E4 recordings from 16 participants aggregated to minute-level HR. In the example shown for Subject 12 on 2020-05-04, HRPI was 105, indicating at least 105 minutes with HR ≥105 bpm (Fig. 1B). Because HRPI is defined from the full ranked distribution, it is threshold-free and nonparametric, jointly capturing the intensity and persistence of HR elevation without relying on an arbitrary predefined cutoff.

### HRPI captures sustained high HR beyond mean HR

To determine what aspect of daily HR behavior is captured by HRPI, we examined its relationships with conventional HR metrics across subject-days. Mean HR was included as the most direct summary of overall HR level, whereas the coefficient of variation (CV) of HR was included to reflect dispersion of HR values within a day. HRPI was positively associated with mean HR (*r =* 0.83) and also with CV HR (*r =* 0.48), indicating that HRPI is related both to overall HR level and to the distribution of HR values (Fig. 2A,B).

**Fig. 2.**
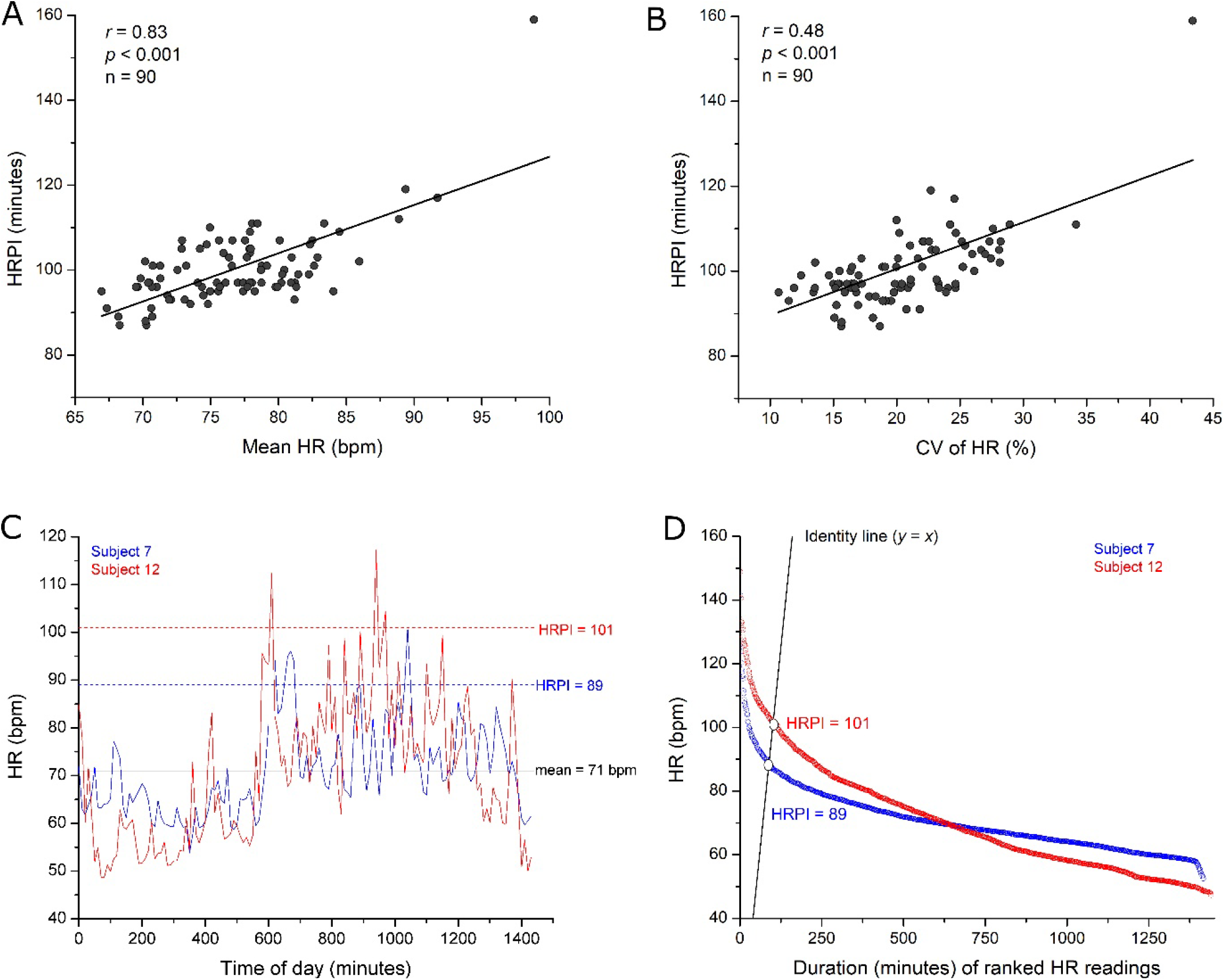
Association of HRPI with HR level and variability. A) HRPI is correlated with mean HR (*r =* 0.83) and B) coefficient of variation (CV) of HR (*r =* 0.48). C-D) Two representative days with similar mean HR but different HRPI: Subject 007, 2020-03-18 (mean HR 71 bpm; HRPI 89) and Subject 012, 2020-05-04 (mean HR 71 bpm; HRPI 101). Minute-level HR is shown as 10-minute averages for visualization. The higher-HRPI day (Subject 012) shows more sustained elevated HR.

Despite the strong correlation between HRPI and mean HR, heterogeneity was evident among days with similar mean HR. As an example, two representative subject-days each had a mean HR of 71 bpm but different HRPI values: Subject 007 on 2020-03-18 had an HRPI of 89, meaning at least 89 minutes with HR ≥89 bpm, whereas Subject 012 on 2020-05-04 had an HRPI of 101, meaning at least 101 minutes with HR ≥101 bpm (Fig. 2C,D). Relative to Subject 007, Subject 012 showed more sustained elevation of HR across the day, consistent with its higher HRPI and indicating that HRPI captures information beyond mean HR alone.

Because HRPI was strongly correlated with mean HR, we next examined whether HRPI contains information beyond average HR level. To test this, we used linear regression to separate the part of HRPI explained by mean HR from the part that remained unexplained. The unexplained part, termed residual HRPI, represents the component of HRPI independent of mean HR. Residual HRPI was no longer associated with mean HR, confirming effective removal of the mean-related component (Fig. 3A). Residual HRPI remained strongly associated with CV HR (*r =* 0.92, *p <* 0.001, n = 90), indicating that HRPI reflects a variability-related component in addition to overall HR level (Fig. 3B).

**Fig. 3.**
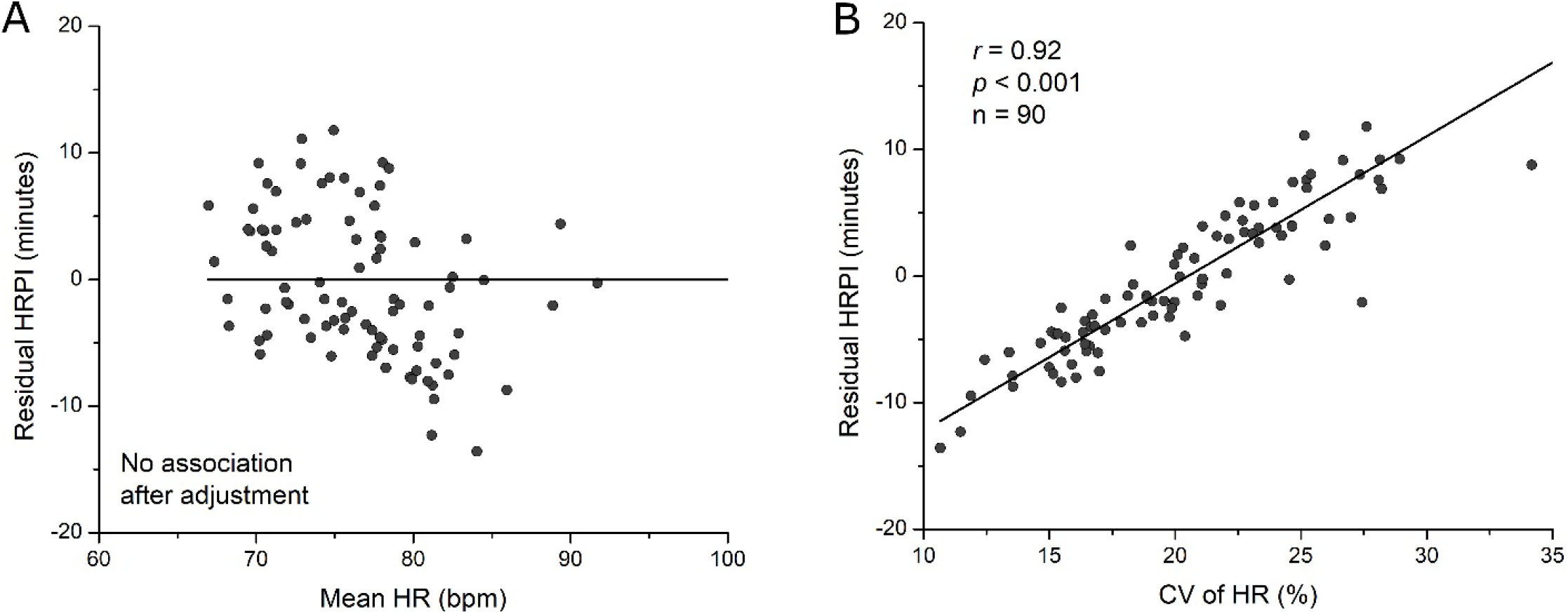
Residual HRPI reveals structure beyond mean HR. (A) Residual HRPI (adjusted for mean HR) shows no association with mean HR, confirming removal of the mean component. (B) Residual HRPI is strongly associated with HR variability (CV) (*r =* 0.92, *p <* 0.001, n = 90), indicating that HRPI captures variability-related structure beyond mean HR.

### HRPI shows day-to-day stability within individuals

For a daily HR metric to be useful in longitudinal monitoring, it should show reasonable reproducibility across nearby days within the same individual, so that changes in the metric are more likely to reflect meaningful physiological differences than excessive day-to-day noise. The PhysioNet “BIG IDEAs Glycemic Variability and Wearable Device” dataset is well suited for this purpose because it provides repeated daily wearable measurements from 16 participants over an approximately two-week period, enabling assessment of within-subject day-to-day stability.

To place HRPI in context, we compared it with conventional HR metrics that represent complementary features of the daily HR profile: mean HR as a measure of overall HR level; p90 HR and p95 HR as summaries of the upper tail of the HR distribution; CV HR and SD HR as measures of day-level HR variability; and minutes with HR >100 bpm as a threshold-based measure of tachycardia burden. Stability was quantified as the within-subject coefficient of variation (CV) across days.

Across individuals, mean HR showed the greatest day-to-day stability, with a median within-subject CV of 4.67%. HRPI was nearly as stable, with a median within-subject CV of 5.49%, and showed stability comparable to the upper-tail metrics p90 HR (6.38%) and p95 HR (6.48%). In contrast, variability-based metrics were substantially less stable, with median within-subject CVs of 13.21% for CV HR and 15.52% for SD HR. The threshold-based metric minutes with HR >100 bpm was by far the least stable, with a median within-subject CV of 33.33% (Fig. 4). Together, these findings indicate that HRPI preserves favorable day-to-day stability within individuals while remaining more robust than variability-based and threshold-based summaries of daily HR.

**Fig. 4.**
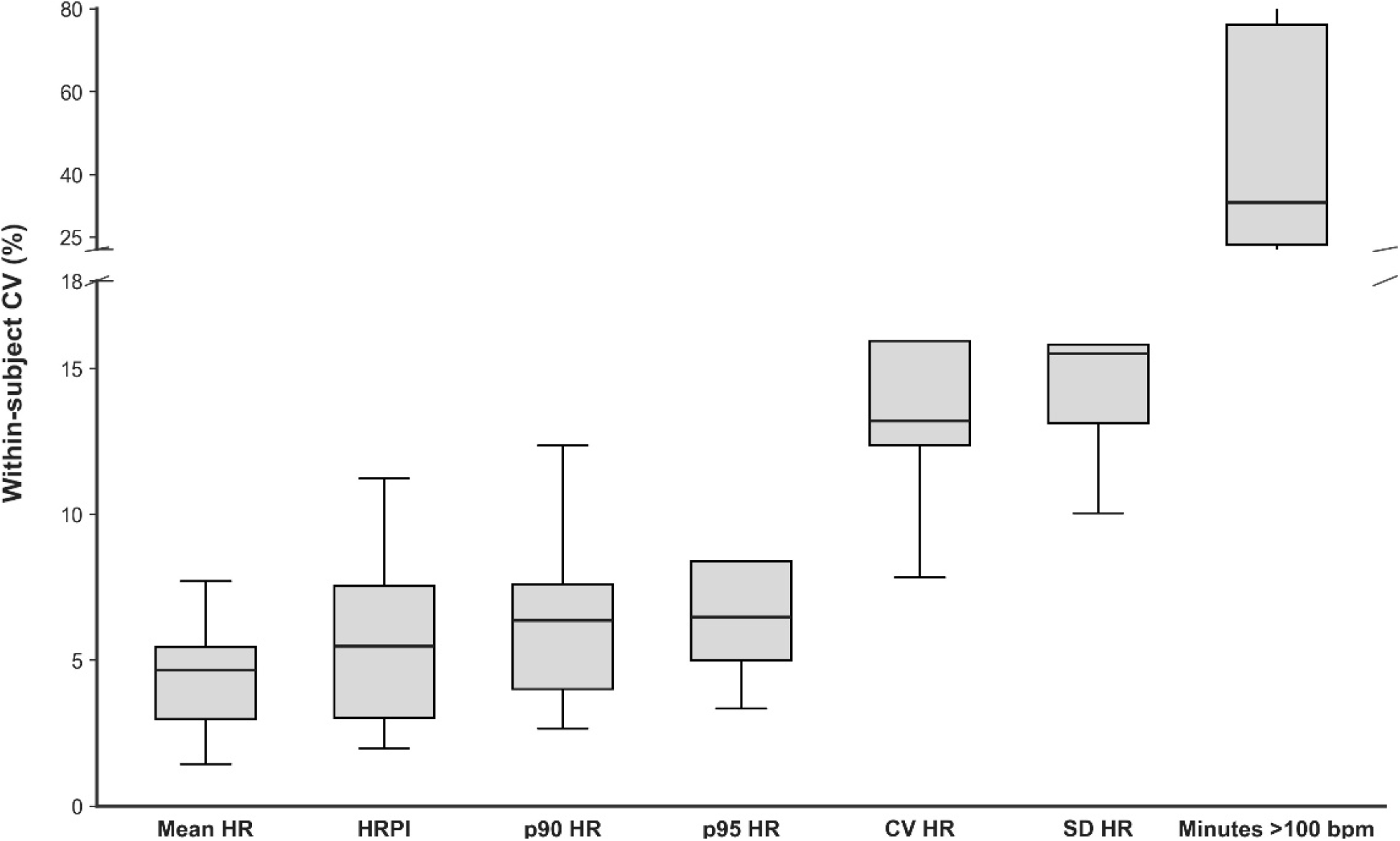
Within-subject stability across HR metrics. Boxplots show median and interquartile range (IQR) of within-subject coefficient of variation (CV) across subjects for Mean HR, HRPI, p90 HR, p95 HR, CV HR, SD HR, and Minutes >100 bpm. Whiskers extend to 1.5×IQR with outliers suppressed. For some metrics (e.g., p95 HR), upper whiskers coincide with the box due to a tightly distributed upper range. A broken y-axis is used to accommodate the higher variability of Minutes >100 bpm. HRPI shows stability comparable to mean HR and percentile summaries and greater stability than variability and threshold-based metrics.

### HRPI decreases with age in healthy subjects

Because HR is strongly age dependent in healthy populations, we next examined whether HRPI shows a corresponding age relationship in an independent healthy reference cohort. For this analysis, we used a subject-level dataset derived from the healthy RR interval cohort, including subject ID, age, and HRPI values[8]. The table initially contained 147 rows; after exclusion of subjects with missing age or HRPI, 138 subjects remained for analysis, spanning infancy through adulthood. Because the cohort was concentrated at younger ages, age was displayed on a logarithmic scale in the figure to improve visualization, whereas regression analyses were performed on the raw age values.

HRPI showed a strong inverse association with age across the cohort. Pearson correlation demonstrated a significant negative relationship between HRPI and age (*r =* -0.715, *p* = 7.273 × 10^-23^), and linear regression showed a slope of -1.657 HRPI units per year with an R^2 of 0.511 (Fig. 5). Thus, older age was associated with lower HRPI values, and age explained approximately half of the between-subject variation in HRPI in this healthy dataset. These findings indicate that HRPI declines progressively with age in healthy subjects, consistent with the known age dependence of HR[9].

**Fig. 5.**
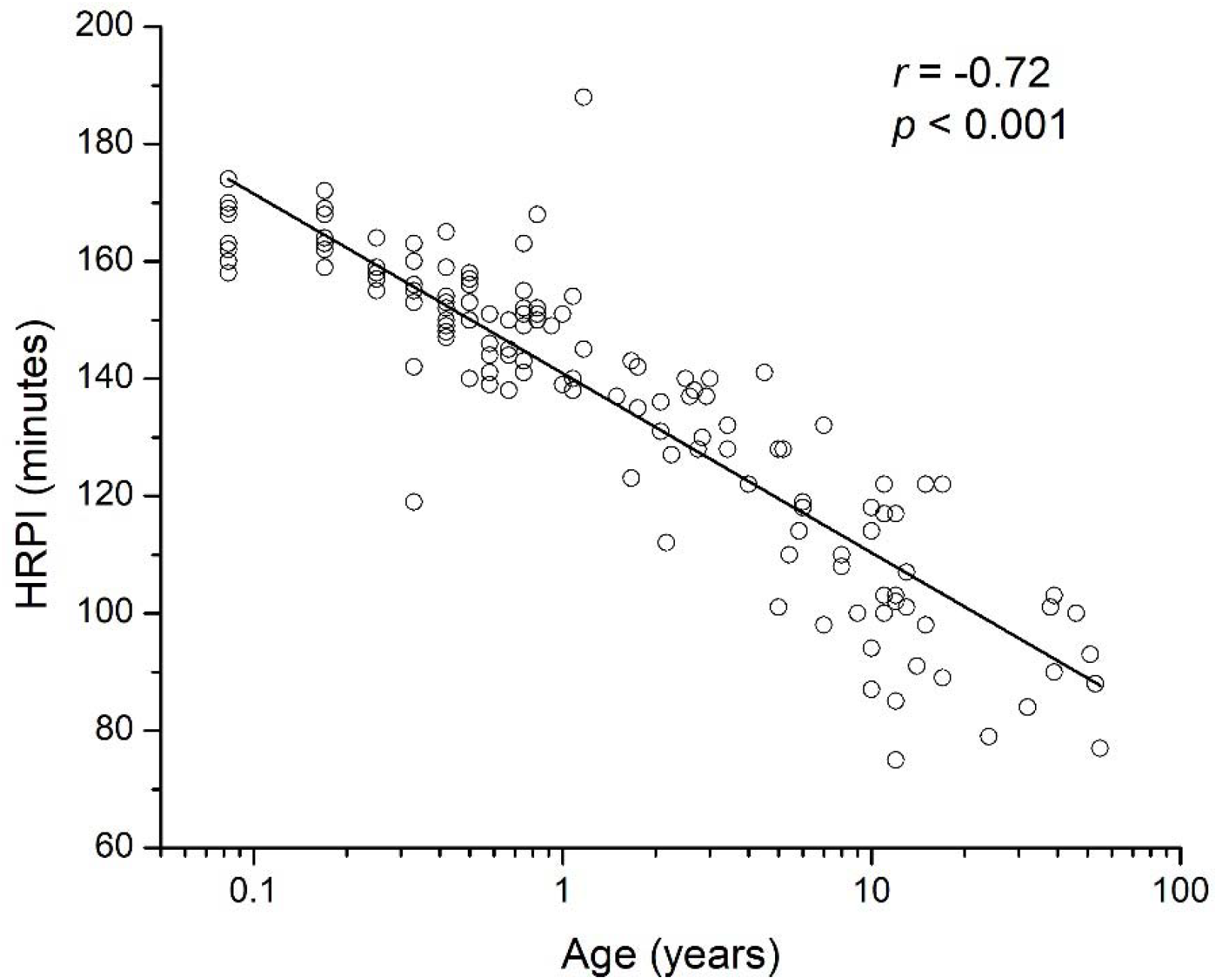
Relationship between HRPI and age. Each point represents an individual subject from the PhysioNet healthy RR interval dataset. Age is shown on a logarithmic scale for visualization. The solid line indicates a linear regression fit (*r =* −0.72, *p <* 0.001).

## Discussion

In this study, we proposed HRPI and found that it was closely related to conventional HR metrics, yet was not fully explained by mean HR alone. HRPI also showed relatively low day-to-day variation within individuals. As expected, HRPI was strongly correlated with mean HR, indicating that overall HR level is a major determinant of the metric. However, days with identical mean HR could still show different HRPI values, and regression-based adjustment further showed that the component of HRPI not explained by mean HR remained strongly associated with HR variability. Together, these findings suggest that HRPI reflects information related to daily HR structure beyond mean HR alone. In addition, HRPI showed low within-subject day-to-day variation, with stability close to mean HR and percentile-based summaries and clearly better than variability-based metrics and the threshold-based metric minutes with HR >100 bpm. The inverse association between HRPI and age in healthy subjects further supports the physiological relevance of the metric.

Together, these observations suggest that HRPI may serve as a simple daily summary of sustained HR elevation. HRPI is threshold-free, and readily interpretable, and it combines two complementary dimensions of the daily HR profile, magnitude and duration, into a single quantity. Unlike maximum HR, it is not driven by brief peaks, and unlike minutes above a fixed threshold, it does not depend on an arbitrary cutoff. Compared with percentile-based metrics, HRPI has a more direct interpretation because its value can be translated back into time spent at or above the same HR level. For example, an HRPI of 100 indicates at least 100 minutes with HR ≥100 bpm. Another practical feature is that the metric is not inherently tied to a specific acquisition platform. In this study, HRPI was derived both from wearable HR recordings and from Holter-derived RR interval data converted to HR, suggesting that the concept is portable across monitoring modalities as long as comparable HR time series are available.

Several limitations should be noted. The wearable analysis was based on a modest sample size, and the stability analysis was performed in a relatively short repeated-measures window of about two weeks. The wearable and healthy reference cohorts also differed in population characteristics and signal source, with one based on wearable HR recordings and the other on RR interval data converted to HR. In addition, the age analysis was cross-sectional and was included mainly to support physiological relevance rather than to establish a new age-dependent phenomenon. Finally, the present study did not test HRPI against clinical outcomes or disease phenotypes. Future studies should evaluate HRPI in larger and more diverse cohorts, determine its relationship to clinically relevant phenotypes such as elevated resting HR and tachycardia burden, and assess its value in longitudinal monitoring, exercise and recovery physiology, autonomic phenotyping, and outcome-oriented clinical studies.

## Conclusion

In summary, HRPI provides a simple and readily-interpretable daily summary of sustained HR elevation that integrates both magnitude and duration without requiring an arbitrary threshold. HRPI showed robust day-to-day stability within individuals. These features support HRPI as a practical candidate for summarizing persistent elevation of HR in longitudinal HR data. Future studies should determine its utility in larger cohorts and in clinically relevant settings. Its intuitive and easily interpretable format may facilitate broad use by both specialists and non-specialists.

## Author Contributions

R.Z. conceptualized, designed the study, analyzed the data, and wrote the paper.

## Data Availability Statement

The heart rate dataset analyzed in this study is publicly available from PhysioNet. The Python scripts used for analysis are available from the corresponding author upon reasonable request.

